# Auditory Stimulus Information Entropy Modulates Inter-Brain Synchronization: Evidence from Wireless EEG Hyperscanning

**DOI:** 10.1101/2025.10.14.681560

**Authors:** Junhao Liao, Gan Huang, Wanru Zhao, Cindy Xindi Li, Pak Wing Calvin Cheng, Rui Sun, Hsiang-Yu Yuan, Junling Gao, Rainbow Tin Hung Ho

**Affiliations:** Centre on Behavioral Health, Faculty of Social Sciences, The University of Hong Kong, Pokfulam, Hong Kong SAR, China; The School of Biomedical Engineering, Medical School, Shenzhen University, Shenzhen, China; The Guangdong Provincial Key Laboratory of Biomedical Measurements and Ultrasound Imaging, Shenzhen, China; Department of Social Work and Social Administration, Faculty of Social Sciences, The University of Hong Kong, Pokfulam, Hong Kong SAR, China; Department of Psychiatry, The University of Hong Kong, Hong Kong, China; Department of Rehabilitation Sciences, The Hong Kong Polytechnic University, Hong Kong SAR, China; Department of Biomedical Sciences, Jockey Club College of Veterinary Medicine and Life Sciences, City University of Hong Kong, Hong Kong Special Administrative Region of China; Centre of Buddhist Studies, The University of Hong Kong, Hong Kong SAR, China

**Keywords:** Multi-person EEG hyperscanning, Information entropy, Inter-brain synchronization, Phase-locking value, Neural oscillations

## Abstract

Inter-brain synchronization (IBS) – reflecting inter-individual correlated neural activity during interaction – marks shared experiences like music listening. The ability of complex auditory stimuli (e.g., music) to induce IBS links to their information dynamics, notably the uncertainty they evoke, which challenges the nervous system’s predictive coding. Based on mutual prediction theory (interacting individuals simultaneously process their own behavior and predict their partner’s; accurate mutual predictions lead to convergent neural representations and thus IBS), this study hypothesized that higher stimulus uncertainty enhances IBS (heightened uncertainty reduces independent predictability, promoting convergent representations and stronger IBS). Using information entropy to quantify uncertainty, the study conducted hyperscanning, manipulated entropy across Resting State and 6 Hz auditory stimuli (ASSR, MMN, AHER, Dream Wedding), and measured IBS via phase-locking values (PLV). Results showed frequency specificity: 6 Hz PLV increased with entropy (DW ≈ AHER > MMN ≈ ASSR > Resting State); Alpha band had highest PLV in Resting State. Critically, PLVs differed significantly between any two conditions, and each experimental condition’s PLV was also significantly different from that of the Resting State. Findings confirm a 6 Hz-specific positive association between auditory uncertainty and IBS, suggesting musical elements may facilitate social interaction by modulating entropy, with entropy-IBS relations showing frequency dependence.

## INTRODUCTION

The human capacity for complex social interaction is a fundamental aspect of our species, relying on sophisticated cognitive processes (Arioli et al., 2018; Frith & Frith, 2012). Social cognitive neuroscience has made significant progress in exploring the mechanisms by which individual brains process social information and engage in interpersonal interactions (Hari & Kujala, 2009; Van Overwalle, 2008). However, social interactions involve complex nonlinear systems. These systems arise from the collaborative process of two or more individuals, and their underlying mechanisms cannot be deduced by simply summing up effects observed in isolated brains (Hasson et al., 2012; Koike et al., 2015; Konvalinka & Roepstorff, 2012). To address this, hyperscanning technology, that enables synchronously record brain activity signals from multiple individuals and quantitatively analyze inter-brain synchronization (IBS), has emerged as a pivotal methodology (Babiloni & Astolfi, 2014; Czeszumski et al., 2020). IBS has been proposed as a neural correlate of social alignment and is increasingly recognized as a key mechanism facilitating interpersonal coordination and shared understanding (Gvirts & Perlmutter, 2020).

IBS arises from shared cognitive processes during social interaction (Zheng et al., 2020). The co-representation mechanism establishes common neural representations of external stimuli, actions, or mental states between individuals (Hasson et al., 2012; Jiang et al., 2021; Shamay-Tsoory et al., 2019), while mutual prediction enables anticipatory coding of partners’ behaviors (Hamilton, 2021). Key neural substrates include the mirror neuron system for action alignment (Lu et al., 2023; Ménoret et al., 2014) and the mentalizing system for inferring intentions (Begliomini et al., 2017; Van Overwalle & Baetens, 2009). To quantify IBS, phase-locking value (PLV) is widely employed, particularly for electrophysiological data (e.g., EEG). PLV measures the consistency of phase differences between two neural signals over time, ranging from 0 (no synchrony) to 1 (perfect phase locking) (Hakim et al., 2023). It isolates phase alignment independent of signal amplitude, making it ideal for detecting oscillatory coupling in specific frequency bands (Hakim et al., 2023).

Previous research indicates that music listening serves as an effective auditory stimulus for inducing IBS during dyadic interactions (Cheng et al., 2024). Music listening constitutes a shared experience that promotes inter-subject synchronization of neural activity, facilitating emotional and cognitive alignment (Sihvonen & Särkämö, 2022). Hyperscanning research has demonstrated that inter-brain synchronization (IBS) during music perception arises through three key mechanisms: (1) shared auditory processing (Lindenberger et al., 2009), (2) emotional contagion (Chabin et al., 2022), and (3) predictive timing and attention (Szymanski et al., 2017). Empirical studies employing neural recordings and controlled auditory stimuli have revealed significant stimulus-response correlations and inter-subject neural coupling during naturalistic music listening (Kaneshiro et al., 2020).

This synchronization occurs across multiple brain regions, including the auditory midbrain, thalamus, auditory cortex, frontal and parietal cortices, and motor planning areas, with more pronounced effects observed for natural music than pseudo-musical stimuli (Abrams et al., 2013). Notably, natural music elicits stronger IBS compared to artificially constructed pseudo-musical stimuli (Kaneshiro et al., 2020). Furthermore, IBS is modulated by stimulus familiarity, with repeated exposure to the same music leading to decreased neural coupling (Dauer et al., 2021; Madsen et al., 2019). A similar pattern of dynamic functional network synchronization has been observed in preterm infants during naturalistic music listening, particularly involving the right temporal, frontal, and motor cortices (Ren et al., 2022).

Emerging evidence suggests that IBS may correlate with subjective experiential states. Stronger synchronization has been reported when listeners concurrently experience similar levels of musical pleasure (Chabin et al., 2022; Ueno & Shimada, 2023). While the functional implications of music-induced IBS remain underexplored, preliminary findings suggest a potential role in enhancing group creativity (Hosseini et al., 2019).

The ability of complex auditory stimuli such as music to elicit IBS is tied to their information dynamics— specifically, the uncertainty they introduce into neural prediction systems (Gold et al., 2019; Lumaca et al., 2019; Pearce, 2018; Pearce & Wiggins, 2012). Music exemplifies a high-uncertainty auditory environment due to its dynamic variations in rhythm, harmony, and spectral features (Gold et al., 2019). To quantify such uncertainty across diverse auditory inputs, this study employ information entropy—a universal metric capturing the unpredictability or disorder within any signal sequence (Hansen & Pearce, 2014; Pearce & Wiggins, 2012; Shannon, n.d.). The higher the entropy, the more unpredictable the signal is in its temporal or frequency structure and the richer its dynamic variations; the lower the entropy, the more it manifests as redundant, predictable regular structures (Shannon, n.d.).

In the mutual prediction theory, during social interactions, individuals concurrently process their own behavioral information (A_self_/B_self_) and predicted behavioral information of others (A_other_/B_other_). When predictions are accurate, their neural representations (A_self_ + A_other_ ≈ B_self_ + B_other_) converge, resulting in IBS (Hamilton, 2021; Kingsbury et al., 2019). Therefore, according to this theory, it is speculated that the information entropy of the stimulus may influence IBS by affecting individuals’ processing of their own behavioral information and their prediction of others’ behavior through uncertainty.

Although existing research using paradigms such as Auditory Steady-State Response (ASSR) and Mismatch Negativity (MMN) has revealed how individual brains process uncertainty (Liang et al., 2023; Lumaca et al., 2019), few studies have extended this mechanism to the group level, particularly lacking systematic investigation into how auditory stimulus uncertainty modulates IBS. Concurrently, while extensive literature has explored the impact of musical attributes (rhythm, melody, emotion) or social contexts on IBS (Cheng et al., 2024), scant attention has been given to systematically evaluating the direct regulatory effects of information-theoretic properties—especially information entropy—of auditory stimuli on IBS. Therefore, this study focuses on auditory stimuli with varying entropy levels, including high-entropy musical conditions, aiming to address these gaps by examining how stimulus entropy drives IBS in multi-person groups.

To manipulate the information entropy levels of auditory stimuli, this study employed three classical paradigms—Auditory Steady-State Response (ASSR), Mismatch Negativity (MMN), and Auditory High Entropy Response (AHER). Research by Huang and co-workers (2023) demonstrates their ascending entropy gradient (ASSR = 0, MMN = 0.29, AHER = 1), fulfilling the present study’s hierarchical entropy manipulation requirements (Liang et al., 2023). Moreover, given music’s inherent structural complexity (Toader et al., 2023), the present study incorporated Dream Wedding (DW; entropy = 2.07) to extend the entropy spectrum and probe music’s entropy-mediated impact on IBS. Consequently, five conditions were implemented: Resting State, ASSR, MMN, AHER, and DW—systematically comparing auditory entropy levels (0–2.07) on group neural synchronization.

## METHODS

### Participants

Participants in this study were recruited based on the following inclusion criteria: (1) age between 18 and 50 years; (2) native speaker of either Cantonese or Mandarin; (3) right-handed; (4) no professional music training within the past three years; (5) no history of diagnosed psychiatric disorders (e.g., anxiety, depression, schizophrenia); (6) no use of anesthetics, sedatives, or stimulants within the past year; (7) no history of brain injury, brain surgery, stroke, or epilepsy; and (8) a regular sleep schedule with at least 7 hours of sleep per night during the three days preceding the experiment.

A total of four healthy, right-handed female participants (age range: 18-30 years) who met all the above criteria were included in this study.

Ethical approval for this study was obtained from the Institutional Review Board of the University of Hong Kong (EA230384). All participants provided written informed consent prior to their inclusion.

### Procedure

The experiment comprised five eyes-closed conditions. Participants first completed a Resting State without auditory stimuli, followed by four randomly ordered auditory conditions: ASSR, MMN, AHER, and DW. Each 3-minute condition was followed by a 20-second interval for completing the Self-Assessment Manikin questionnaire (Betella & Verschure, 2016) on a mobile device, assessing pleasure and arousal responses to stimuli. The entire session lasted approximately 20 minutes.

In view of previous studies demonstrating particularly pronounced effects with 6 Hz stimulation (Liang et al., 2023), all conditions in the present study featured standardized volume and auditory sequences presented at a rate of 6 Hz to facilitate direct comparison with existing research findings. Resting State involves no sound. ASSR delivers a 524 Hz pure tone binaurally. MMN presents 524 Hz and 262 Hz tones at 90% and 10% probability respectively. AHER uses equal probability (50%) of the same tones. DW consists of a ‘Dream Wedding’ segment played on electronic piano at a rate of 6 Hz.

**Figure 1.**
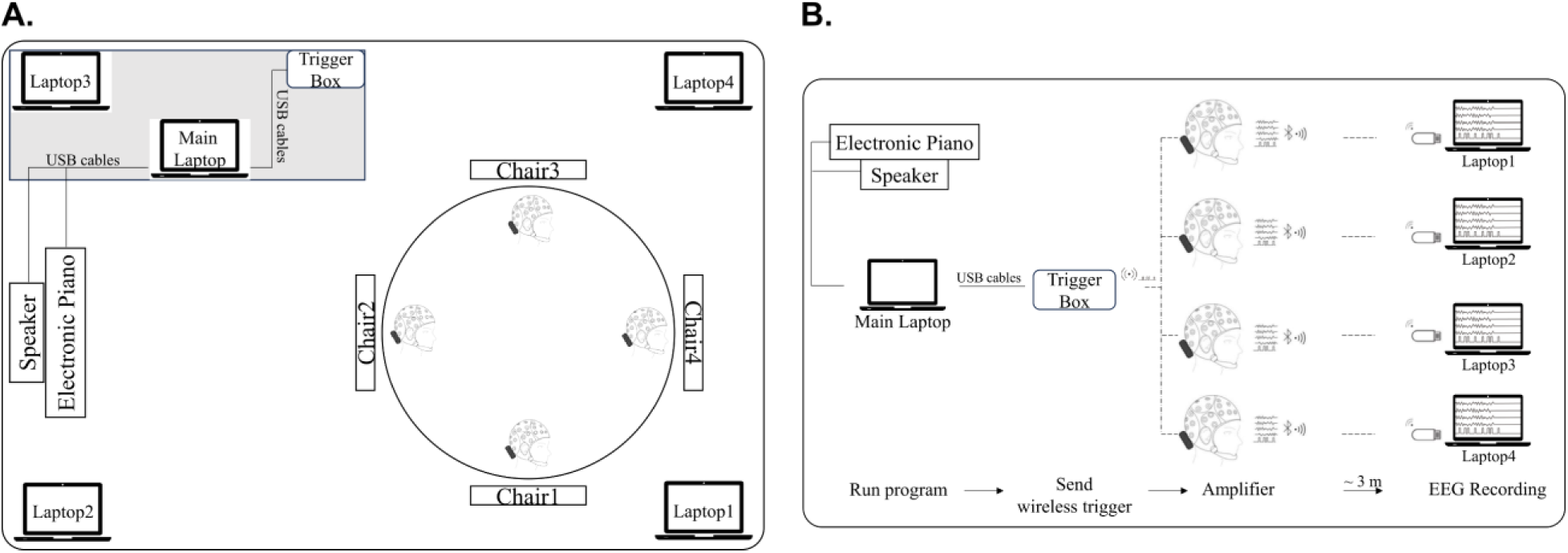
(A) Equipment setup and participant placement. Four participants sat in chairs 1-4. Laptops 1-4 recorded their respective EEG signals. To avoid confusion in EEG data transmission, the current study placed the four laptops used for recording data in the four corners of the room, ensuring that there was a certain distance between each laptop. The main laptop operated the speaker (for ASSR, MMN, AHER stimuli) and the electronic piano (for DW stimuli). The speaker and electronic piano were positioned together and volume-matched. (B) Hyperscanning information flow schematic diagram.

### Data Acquisition

This four-person EEG hyperscanning study was conducted in an electromagnetically shielded quiet room. Prior to EEG recording, participants washed their hair thoroughly and ensured adequate sleep the night before testing. On the experimental day, individuals refrained from neuromodulation therapies (e.g., TMS/tDCS) to prevent EEG interference. During the session, participants maintained comfortable seated positions while minimizing head or body movements.

EEG data acquisition employed a wireless 16-channel EEG amplifier (Shanghai idea-interaction Tech. Co., Ltd., Shanghai, China) with electrodes arranged in the 10-20 system. Sampling occurred at 500 Hz with ground at AFz and reference at FCz. The O2 electrode recorded cardiac signals below the left clavicle, resulting in 15 scalp electrodes plus ECG. All non-ECG electrodes maintained impedance below 30 kΩ. The triggers are wirelessly sent to the four amplifiers simultaneously via the same trigger box, ensuring precise EEG signal synchronization during offline analysis.

### Analysis and Statistics

Analyses used MATLAB with the EEGLAB (Delorme & Makeig, 2004), FieldTrip (Oostenveld et al., 2011), and Letswave7 toolboxes for preprocessing, ERP extraction, time-frequency decomposition, and visualization. EEG preprocessing involved ECG channel removal (unanalyzed), 0.5-30 Hz FIR bandpass filtering, epoch extraction/merging per condition, ICA-based ocular artifact removal, average mastoid rereferencing (TP9/TP10), and condition-based epoch separation.

EEG epochs were extracted from −500 ms to 1000 ms relative to condition-specific markers: standard (*S*) and deviant (*D*) stimuli in ASSR/MMN/AHER conditions, and individual musical note onsets in the DW condition. The segmented epochs were then averaged across trials to derive event-related potential (ERP) components. Peak-to-peak amplitudes were computed as the difference between maximum and minimum voltage values within the 50–250 ms post-stimulus window of ERP waveforms. Compared with the conventional ERP analysis, the Stimulus Onset Asynchrony of 6 Hz stimulation (SOA=166.67 ms) in this work was shorter than the epoch window (−500–1000 ms). The overlap between the two successive stimuli existed commonly for the time domain analysis. Hence, the pre-stimulus ERP is also informative, which could not be treated as the conventional baseline. Hence, this study did not perform baseline correction in the time domain analysis. After the pre-processing of 0.5-30Hz bandpass filtering, the DC component has been effectively removed.

Spectral power was obtained by performing Fast Fourier Transform (FFT) on EEG segments using the Letswave7 toolbox.

Inter-brain synchronization was computed via Hilbert-transform-based bandpass filtering across delta (1–4 Hz), theta (4–8 Hz), alpha (8–14 Hz), and beta (14–30 Hz) bands, extracting instantaneous phases for Phase Locking Value (PLV) calculation. For 6 Hz signals, wavelet transforms derived phase information. PLV (Müller et al., 2013) was defined as:

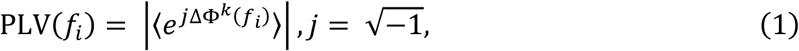

where 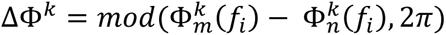 is the phase difference at the central frequency *f_i_* between the instantaneous phases of the two signals m and n across k data points in the segment; 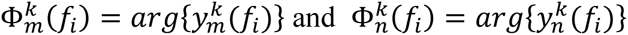. Using 2000-ms sliding windows (200-ms step), PLV was computed for all pairwise electrode combinations across participants (14 electrodes × 4 subjects; total pairs = 56^2^). Results specifically focused on the FCz-FCz electrode pair. The sliding window procedure yielded 891 condition-specific PLV time windows.

Data from four participants prompted exploratory descriptive statistics for ERP peak-to-peak amplitudes and spectral power. For inter-brain PLV analysis, multilevel mixed-effects modeling addressed three nested levels: time windows (891/condition), subject pairs (6 dyads from 4 participants), and conditions (Resting State/ASSR/MMN/AHER/DW). The core model incorporated Condition and Band as fixed effects with random intercepts for pairs and windows. Significant Condition×Band interactions (Satterthwaite-corrected F-tests) prompted band-specific pairwise comparisons with FDR correction. Residual diagnostics validated model assumptions. Analyses used R (4.5.1) with lme4 (1.1-37), lmerTest (3.1-3), and emmeans (1.11.1).

## RESULTS

### Temporal Dynamics of ERP

The grand-averaged ERP responses for ASSR(*S*), MMN(*S*), MMN(*D*), AHER(*S*), and AHER(*D*) are shown in Figure 2(A-C). For the standard stimuli (*S*), the peak-to-peak amplitudes of ASSR, MMN, and AHER are 2.31, 2.37, 2.82 μV, respectively. There appears to be a trend indicating that AHER is larger than MMN and also larger than ASSR.

**Figure 2.**
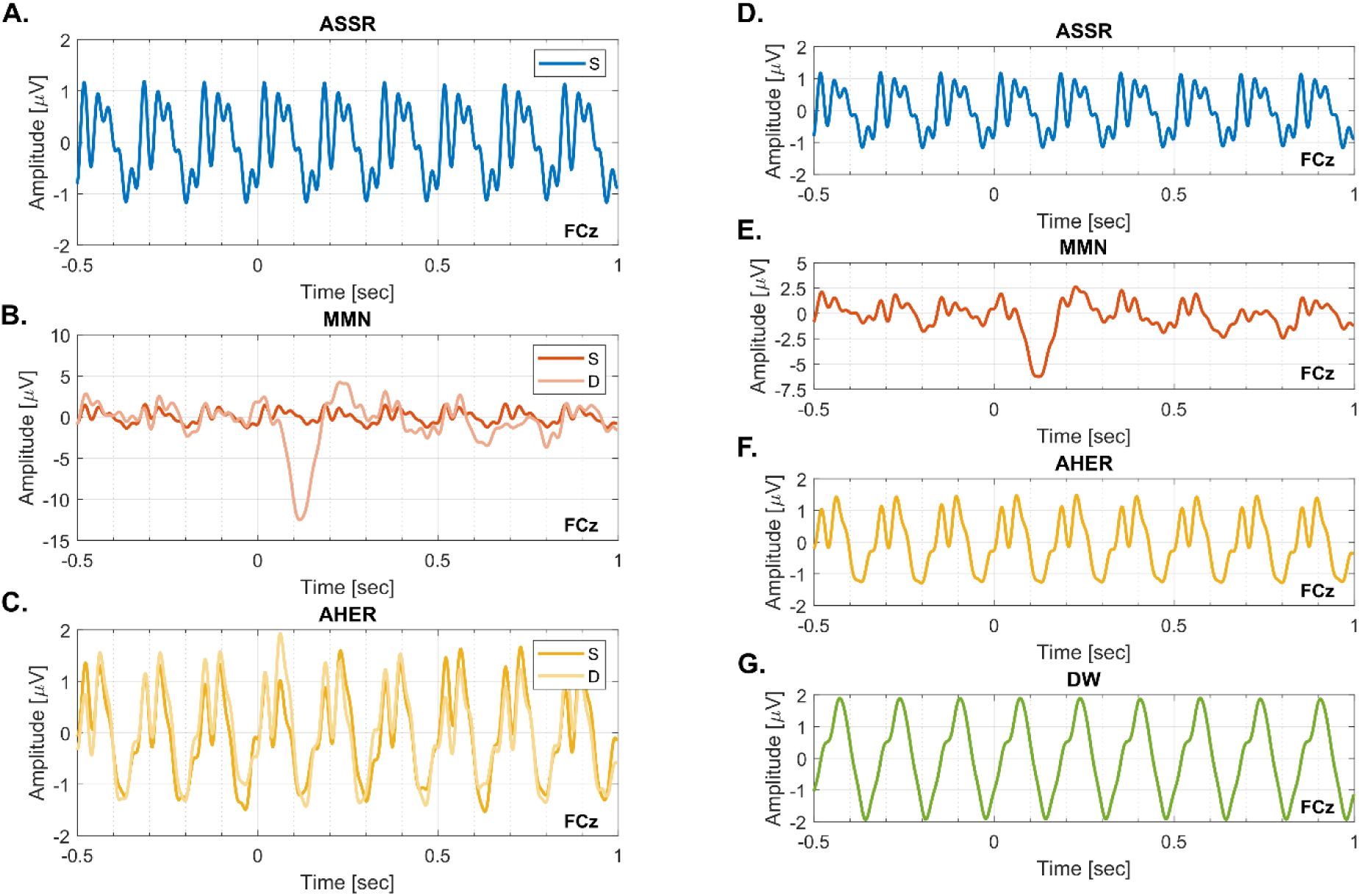
Grand-averaged ERP responses under 6 Hz stimulation at channel FCz. (A-C) ERP responses for ASSR(*S*), MMN(*S*), MMN(*D*), AHER(*S*), and AHER(*D*). “*S*” and “*D*” indicate standard and deviant stimuli. (D-G) ERP responses for ASSR, MMN, AHER, and DW.

For the deviant stimuli (*D*), the peak-to-peak amplitudes of MMN and AHER are 16.71 μV and 3.36 μV, respectively. Therefore, for MMN, the peak-to-peak amplitude for deviant stimuli (16.71 μV) is larger than the peak-to-peak amplitude for standard stimuli (2.37 μV). However, for AHER, the difference in peak-to-peak amplitudes between deviant (3.36 μV) and standard (2.82 μV) stimuli is not obvious.

The grand-averaged ERP responses for ASSR, MMN, AHER, and DW are shown in Figure 2(D-G), with their averaged peak-to-peak amplitudes being 2.31, 8.87, 2.74, and 3.82 μV, respectively. The peak-to-peak amplitudes, from largest to smallest, are ranked as MMN > DW >AHER > ASSR.

### Spectral Characteristics: Power Spectral Density and Power

Fig. 3A presents the power spectral density (PSD) of Resting State (without 6 Hz stimulation) and experimental conditions (with 6 Hz stimulation), demonstrates prominent alpha band activity in the PSD analysis, with a distinct spectral peak observed at 6 Hz outside the alpha frequency range for the experimental conditions. The topographic map shows that the 6 Hz power is concentrated around the FCz electrode.

**Figure 3.**
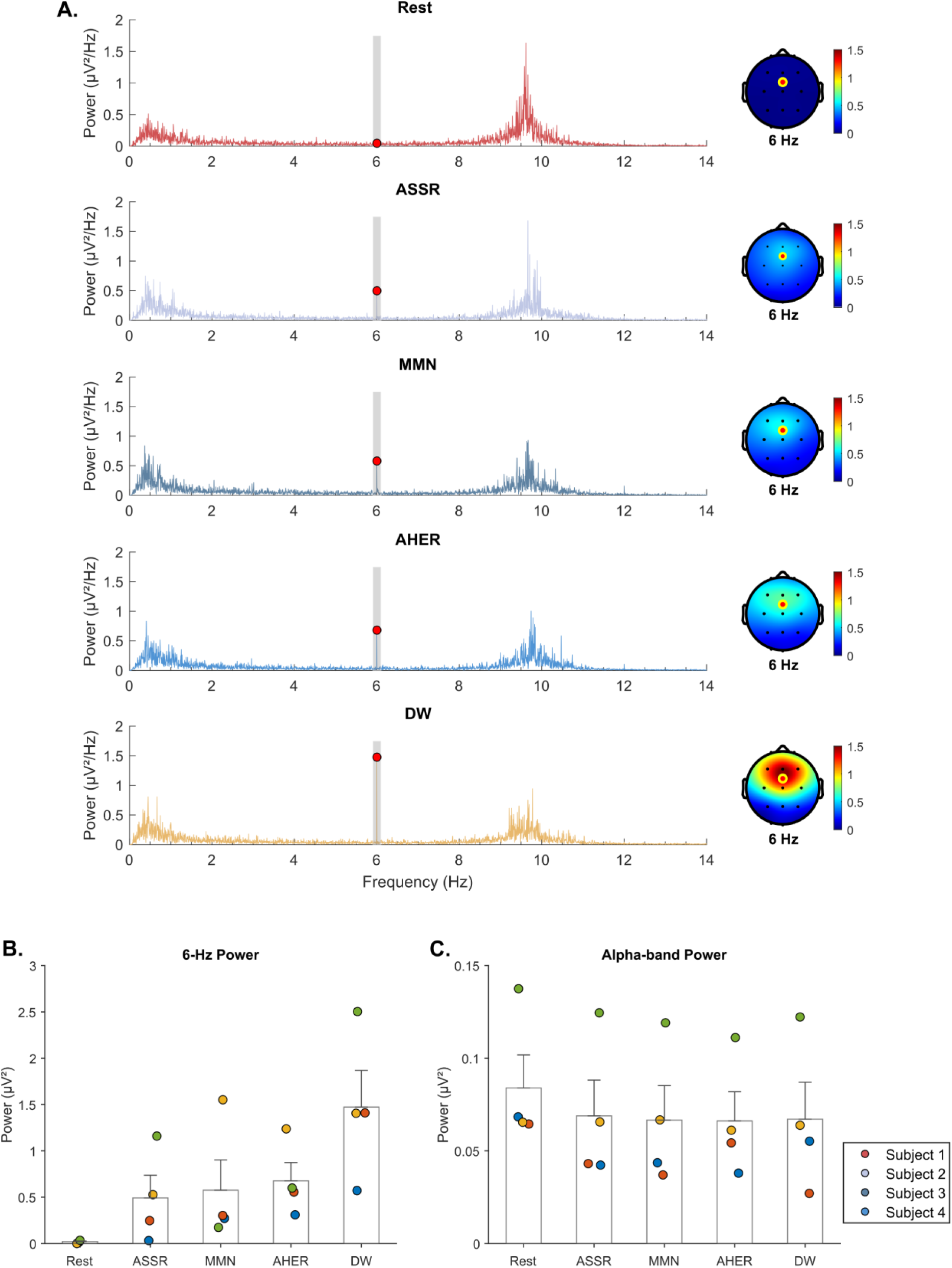
(A) Power spectral density (PSD) for Resting State, ASSR, MMN, AHER, and DW during 6 Hz stimulation at FCz, with the 6 Hz marker indicated on the PSD and channel FCz highlighted on the topographic maps. (B–C) 6 Hz and alpha (8–14 Hz) power values for all conditions at FCz. Dots: individual subjects; bars: mean power; error bars: ±SE.

Fig. 3B and Fig. 3C present individual and mean power values at the 6-Hz band and alpha band (8-14 Hz) across all conditions (Resting State, ASSR, MMN, AHER, DW) for all subjects. For the 6-Hz band, the mean power values of Resting State, ASSR, MMN, AHER, and DW are 0.020, 0.491, 0.574, 0.675, and 1.472 µV², respectively, following the trend DW > AHER > MMN > ASSR > Resting State. For the alpha band, the mean power values are 0.084, 0.069, 0.067, 0.066, and 0.067 µV², respectively. Each experimental condition’s power is less than that of the Resting State.

### Inter-brain Synchronization: Phase Locking Value

The full mixed-effects model revealed significant main effects of Condition (*F*(4, 132730) = 17.877, *p* < 0.001) and Band (*F*(4, 132730) = 9358.180, *p* < 0.001), along with a significant Condition × Band interaction (*F*(16, 132730) = 54.993, *p* < 0.001). These findings indicate that the effects of condition on PLV varied by frequency band. Post-hoc pairwise comparisons, adjusted for false discovery rate (FDR), are detailed below for each band.

In the 6 Hz band (Fig. 4A), Resting State showed significantly lower PLV than ASSR (*β* = −0.019, SE = 0.004, pFDR < 0.001), MMN (*β* = −0.018, SE = 0.004, pFDR < 0.001), AHER (*β* = −0.037, SE = 0.004, pFDR < 0.001) and DW (*β* = −0.042, SE = 0.004, pFDR < 0.001). ASSR showed significantly lower PLV than AHER (*β* = −0.017, SE = 0.004, pFDR < 0.001) and DW (*β* = −0.023, SE = 0.004, pFDR < 0.001). MMN showed significantly lower PLV than AHER (*β* = −0.019, SE = 0.004, pFDR < 0.001) and DW (*β* = −0.024, SE = 0.004, pFDR < 0.001).

**Figure 4.**
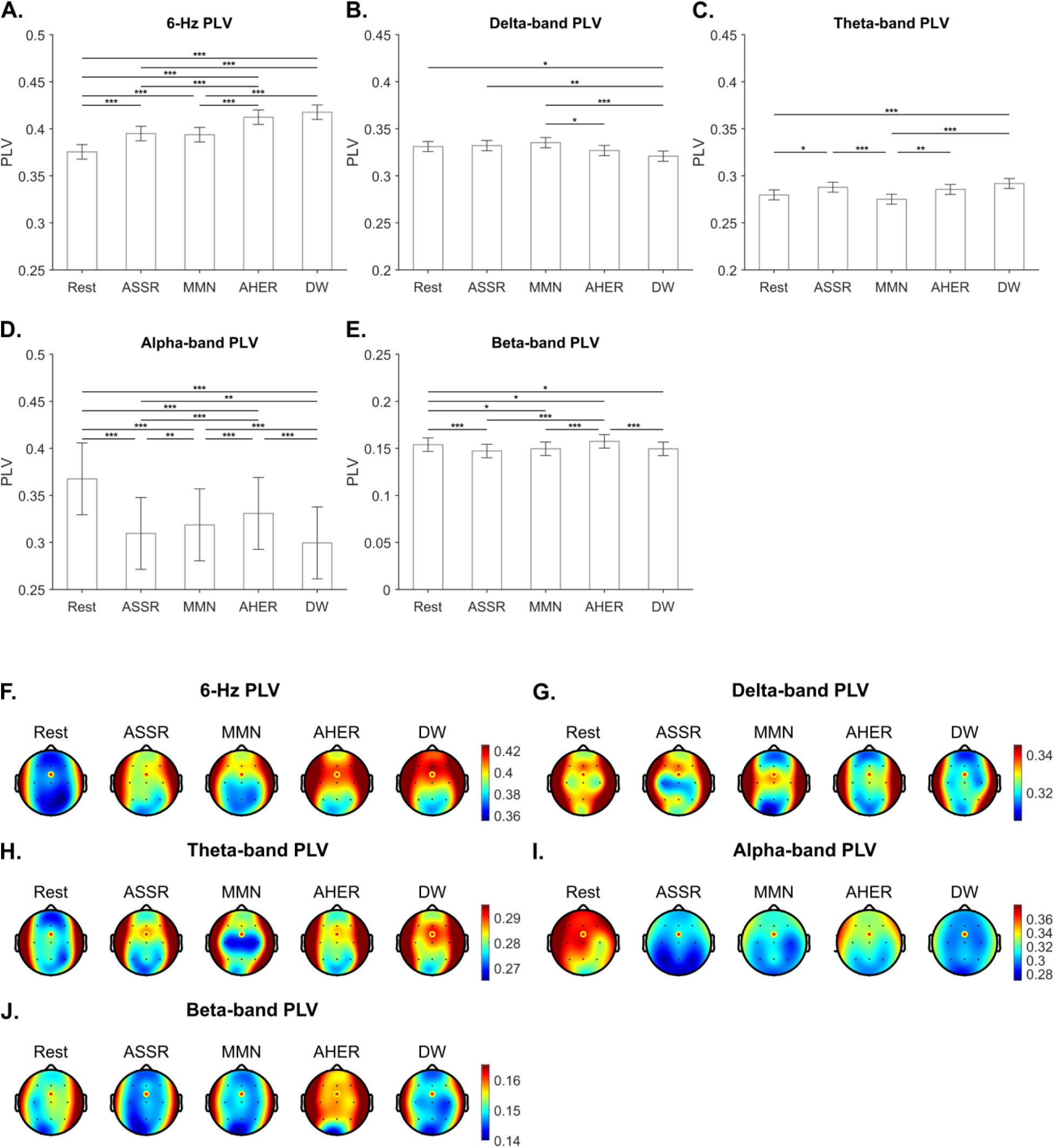
Phase-Locking Value (PLV) at electrode FCz across conditions and frequency bands, and their whole-scalp topographic distributions. (A-E) Mean PLV values at FCz for the Resting State, ASSR, MMN, AHER, and DW conditions in the (A) 6-Hz, (B) Delta, (C) Theta, (D) Alpha, and (E) Beta frequency bands. Error bars represent 95% confidence intervals. Significance levels: *p < 0.05; **p < 0.01; ***p < 0.001. (F-J) Whole-scalp topographic maps illustrate the spatial distribution of PLV for each corresponding frequency band: (F) 6-Hz, (G) Delta, (H) Theta, (I) Alpha, and (J) Beta. The color scale represents PLV magnitude.

In the Delta band (Fig. 4B), Resting State showed significantly higher PLV than DW (*β* = 0.010, SE = 0.003, pFDR = 0.013). ASSR showed significantly higher PLV than DW (*β* = 0.011, SE = 0.003, pFDR = 0.009). MMN showed significantly higher PLV than AHER (*β* = 0.008, SE = 0.003, pFDR = 0.048) and DW (*β* = 0.014, SE = 0.003, pFDR < 0.001).

In the Theta band (Fig. 4C), Resting State showed significantly lower PLV than ASSR (*β* = −0.008, SE = 0.003, pFDR = 0.013) and DW (*β* = −0.012, SE = 0.003, pFDR < 0.001). ASSR showed significantly higher PLV than MMN (*β* = 0.013, SE = 0.003, pFDR < 0.001). MMN showed significantly lower PLV than AHER (*β* = −0.010, SE = 0.003, pFDR = 0.001) and DW (*β* = −0.017, SE = 0.003, pFDR < 0.001).

In the Alpha band (Fig. 4D), Resting State showed significantly higher PLV than ASSR (*β* = 0.058, SE = 0.003, pFDR < 0.001), MMN (*β* = 0.049, SE = 0.003, pFDR < 0.001), AHER (*β* = 0.037, SE = 0.003, pFDR < 0.001), and DW (*β* = 0.068, SE = 0.003, pFDR < 0.001). ASSR showed significantly lower PLV than MMN (*β* = −0.009, SE = 0.003, pFDR = 0.005) and AHER (*β* = −0.021, SE = 0.003, pFDR < 0.001), while showed significantly higher PLV than DW (*β* = 0.010, SE = 0.003, pFDR = 0.002). MMN showed significantly lower PLV than AHER (*β* = −0.012, SE = 0.003, pFDR < 0.001) and significantly higher PLV than DW (*β* = 0.019, SE = 0.003, pFDR < 0.001). AHER showed significantly higher PLV than DW(*β* = 0.031, SE = 0.003, pFDR < 0.001).

In the Beta band (Fig. 4E), Resting State showed significantly higher PLV than ASSR (*β* = 0.007, SE = 0.002, pFDR < 0.001), MMN (*β* = 0.004, SE = 0.002, pFDR = 0.013), and DW (*β* = 0.004, SE = 0.002, pFDR = 0.013), while showed significantly lower PLV than AHER (*β* = −0.004, SE = 0.002, pFDR = 0.036). ASSR showed significantly lower PLV than AHER (*β* = −0.010, SE = 0.002, pFDR < 0.001). MMN showed significantly lower PLV than AHER (*β* = −0.008, SE = 0.002, pFDR < 0.001). AHER showed significantly higher PLV than DW (*β* = 0.008, SE = 0.002, pFDR < 0.001).

## DISCUSSION

This study successfully replicated previous ERP and spectral analysis findings (Liang et al., 2023) and further elucidated the regulatory mechanisms through which stimuli at different information-entropy levels (ASSR, MMN, AHER) and high-entropy music (DW) modulate group-level inter-brain synchrony. The results showed that in the 6 Hz band, PLV increased with information entropy (DW ≈ AHER > MMN ≈ ASSR > Resting State). A similar pattern (AHER > MMN > ASSR) was observed among the ASSR, MMN, and AHER conditions in the alpha band, although the Resting State’s PLV exceeded that of all other conditions.

The ERP analyses revealed three key findings: First, for standard stimuli (*S*), the peak-to-peak amplitude of A-HER significantly exceeded that of both MMN and ASSR. Second, deviant stimuli elicited larger amplitudes than standard stimuli in the MMN paradigm, whereas no significant amplitude difference emerged between deviant and standard stimuli in A-HER. Furthermore, the grand-averaged waveforms demonstrated consistently higher amplitudes for A-HER relative to ASSR. Collectively, these outcomes align directionally with established findings in prior literature (Liang et al., 2023). Spectral analysis revealed that A-HER exhibited significantly greater 6Hz power compared to both ASSR and MMN paradigms. This directional pattern aligns with established findings in prior literature.

For inter-brain synchronization, the results revealed a significant condition-by-frequency interaction. In the 6 Hz band, pairwise comparisons revealed significant differences across all conditions except for ASSR-MMN and AHER-DW contrasts. Notably, the PLV under both AHER and DW conditions significantly exceeded that of the Resting State, ASSR, and MMN conditions. Collectively, inter-brain synchronization strength demonstrated a monotonic increase with information entropy (PLV*high-entropy* > PLV*low-entropy*). Conversely, nonsignificant PLV variations between entropy-matched conditions (e.g., ASSR-MMN vs. AHER-DW) suggest potential stabilization of inter-brain synchrony within specific entropy ranges.

The mutual prediction theory can explain the findings of the present study in the 6 Hz band. Compared with the resting state, low-entropy conditions (ASSR/MMN) enhanced participants’ prediction consistency and thus significantly increased PLV. Likewise, high-entropy conditions (AHER/DW) achieved even greater prediction consistency than the low-entropy conditions, resulting in stronger inter-brain synchronization.

In the alpha band, significant differences in the PLV were observed across all experimental conditions. Notably, the PLV under the ASSR, MMN, AHER, and DW conditions was significantly lower than during the Resting State. This phenomenon may be related to signal-to-noise ratio (SNR) and neuronal synchronous firing (Hülsemann et al., 2019; Lei et al., 2012). Compared to the four 6-Hz auditory stimulation conditions, participants exhibited higher alpha amplitude during the eyes-closed Resting State. This higher amplitude implies a better SNR and more synchronous neuronal firing. Although the calculation of PLV itself does not directly incorporate amplitude, the high and stable alpha amplitude during the Resting State creates essential conditions for measuring relatively higher PLV values by providing a high SNR and intrinsic rhythmic stability. Furthermore, among the three conditions of ASSR, MMN, and AHER, there is a law that PLV increases with the increase of information entropy.

### Limitation and Future Direction

However, it is important to note that this study is limited by its sample size. Future research could address individual differences by increasing the sample size and validating the generalizability of this effect in larger populations. Besides, although the experimental paradigm involved four-person interactions, analytical approaches (e.g., PLV) remained confined to pairwise synchronization assessment (Lee et al., 2019). Future investigations may employ multilayer brain network analysis to advance understanding of collective neurodynamic principles.

## CONCLUSION

This research pioneers the extension of uncertainty processing mechanisms to the group level, systematically elucidating how uncertainty modulates multi-person neural synchrony. Critically, the present study demonstrate that inter-brain synchronization exhibits significant enhancements with increasing information entropy (i.e., heightened uncertainty). This may suggest that the social interaction of listeners can be facilitated by increasing the complexity of the music, specifically in 6 Hz music.

## Supporting information

program code

## ETHICS STATEMENT

The studies involving human participants were reviewed and approved by the Institutional Review Board (IRB), The University of Hong Kong. The participants provided their written informed consent to participate in this study.

## AUTHOR CONTRIBUTIONS

J.L.: Conceptualization, Methodology, Investigation, Formal Analysis, Writing – Original Draft.

G.H.: Conceptualization, Methodology, Resources.

W.Z.: Writing – Original Draft, Writing – Review & Editing.

C.L.: Writing – Review & Editing, Supervision.

P.W.C.C.: Funding Acquisition, Writing – Review & Editing.

R.S.: Funding Acquisition, Writing – Review & Editing.

H.-Y.Y.: Funding Acquisition, Writing – Review & Editing.

J.G.: Conceptualization, Funding Acquisition, Project Administration, Supervision.

R.H.: Conceptualization, Funding Acquisition, Project Administration.

All authors participated in result interpretation, critically reviewed and approved the final manuscript.

## ACKNOWLEDGMENTS

This research is supported by the Seed Fund for Collaborative Research (Grant No. 207080376) at the University of Hong Kong, the National Natural Science Foundation of China (Grant No. 62271326), the Shenzhen Science and Technology Program (Grant No. JCYJ20241202124222027).

## DATA AVAILABILITY STATEMENT

The raw data supporting the conclusions of this report will be made available by the authors, without undue reservation.

