## Supplementary material for "Auditory Stimulus Information Entropy Modulates Inter-Brain Synchronization: Evidence from Wireless EEG Hyperscanning": program code: README.md

**Auditory Stimulus Information Entropy Modulates Inter-Brain Synchronization: Evidence from Wireless EEG Hyperscanning**

This folder contains the following files:

- run_1_resting.m: Script for running during the Resting State.

- run_2_serial.m: Script for running during audio stimulus.

- send_trigger.m: Trigger-sending function called by the first two scripts.

- song2_Trigger.m: Audio trigger function called by the first two scripts.

**Running Steps**

1. First, run run_1_resting.m during the Resting State.

2. Then, run run_2_serial.m during audio stimulus.

Note: send_trigger.m and song2_Trigger.m will be automatically called when the first two scripts are executed and do not need to be run separately.
