## Supplementary material for "Auditory Stimulus Information Entropy Modulates Inter-Brain Synchronization: Evidence from Wireless EEG Hyperscanning": program code: run_1_resting.m

```
% rest: closed eye
clc;clear;

% 静息态时长
rest_time = 180; % unit: second

%% 连接到trigger box，并打开串口
global Obj; % Set global variables
delete(instrfind);
Obj = serialport('COM9', 115200, 'parity', 'none', 'databits', 8, 'stopbits', 1); % Initialize serial port
fopen(Obj);
send_trigger(1, 0);

%% closed
Beeper(1000, 1, 1); % 提示音：音调frequency, 音量fVolume, 长度durationSec
pause(5); % 等待被试平复下来
send_trigger(1, 14);
pause(rest_time); % 静息态阶段
send_trigger(1, 15);
Beeper(1000, 1, 1);

%% 关闭串口
fclose(Obj);
clear global Obj
```
