## Supplementary material for "Auditory Stimulus Information Entropy Modulates Inter-Brain Synchronization: Evidence from Wireless EEG Hyperscanning": program code: run_2_serial.m

```
%% link to trigger box and singlechip
% Elapsed time is 26.266217 seconds.???????????????????
clc;clear;close all;
tic
questionnaire_time = 25;
global play_time
play_time = 180 + 1; % 記得改回180

%% 连接到单片机Arduino
delete(instrfind);
scom = serial('COM6');
fopen(scom);

%% 连接到trigger box，并打开串口
global Obj;
Obj = serialport('COM9', 115200, 'parity', 'none', 'databits', 8, 'stopbits', 1);
fopen(Obj);
send_trigger(1, 0);

%% high frequency entropy
trig1=hex2dec(['01';'55';'66';'0D']); % ASSR 条件的开始
trig2=hex2dec(['02';'55';'66';'0D']); % MMN 条件的开始
trig3=hex2dec(['03';'55';'66';'0D']); % AHER 条件的开始
trig4=hex2dec(['04';'55';'66';'0D']); % note1, deviant(D)
trig5=hex2dec(['05';'55';'66';'0D']); % note2, standard(S)
trig6=hex2dec(['06';'55';'66';'0D']); % Dream wedding 条件的开始
trig7=hex2dec(['07';'55';'66';'0D']); % DW note

trig31 = hex2dec(['1F';'55';'66';'0D']); % ASSR 条件的结束：31
trig32 = hex2dec(['20';'55';'66';'0D']); % MMN 条件的结束：32
trig33 = hex2dec(['21';'55';'66';'0D']); % AHER 条件的结束：33
global trig36
trig36 = hex2dec(['24';'55';'66';'0D']); % Dream wedding 条件的结束:36

fig=figure();
set(fig,'DoubleBuffer','on')
set(fig,'MenuBar','None');
set(fig,'ToolBar','None');
% set(fig,'WindowState','Fullscreen');
set(fig,'Color',[0,0,0])
axes('position',[0,0,1,1])
axis off;
xlim([-1920,1920]/2)
ylim([-1080,1080]/2)
txt1=text(0,0,'+','Fontsize',300,'color','w','HorizontalAlignment','center');
txt2=text(0,0,'*','Fontsize',100,'color','w','HorizontalAlignment','center','visible','off');
rect=rectangle('Position',[-150,-150 300 300],'FaceColor','w','visible','off');
pause(3);

%
% freq=[0.5,1,2,3,4,5,6,7,8,10,12];
% dur=[80,40,20,20,20,60,20,20,20,20,20];

freq=[0.5 6 6]; % frequency, Hz
dur=[80,20,play_time]; % duration time, second
% dur=[8000,400,200,200,20,20,20,20,20,20,20];
% dur=[2000,2000,2000,2000,2000,2000,2000,2000,2000,2000,2000];
prop=[0,0.05,0.5]; % probability, i.e. ASSR, MMN, AHER

rng('shuffle')
for times=1
    for cond_idx = randperm(4) % 1则播放ASSR，2则播放MMN，3则播放MMN，4则播放DW
        set(txt2,'visible','off');
        set(txt1,'visible','on');
        
        Beeper(1000, 1, 1); % audio播放前的提示音
        pause();

        if cond_idx == 4
            disp([times,cond_idx]);
            song2_Trigger;
        else
            for seq_idx = 3 % fixed for 180 seconds
                disp([times,cond_idx]);
                % 发送表示条件开始的trigger
                if cond_idx==1 % 如果当前条件是ASSR，则发送trigger "1"
                    fwrite(Obj,trig1);
                end
                if cond_idx==2% 如果当前条件是MMN，则发送trigger "2"
                    fwrite(Obj,trig2);
                end
                if cond_idx==3% 如果当前条件是AHER，则发送trigger "3"
                    fwrite(Obj,trig3);
                end
                pause(0.01);
                tic;
                t=toc;
                k=1;
                while t < dur(seq_idx) + 2 % when the time is less than 20 seconds
                    if t > k/freq(seq_idx) % 1/6 seconds, 2/6 seconds, 3/6 seconds, 4/6 seconds, ...
                        if rand(1,1) < prop(cond_idx)
                            fwrite(Obj,trig4);
                            send_serial(scom,'2'); % 发送2到单片机Arduino
                        else
                            fwrite(Obj,trig5);
                            send_serial(scom,'1'); % 发送1到单片机Arduino
                        end
                        pause(0.01);
                        k=k+1;
                    end
                    t=toc;
                end
            end
        end
        % 发送表示条件结束的trigger
        if cond_idx==1 % 如果当前条件是ASSR，则发送trigger "31"
            fwrite(Obj,trig31);
        end
        if cond_idx==2% 如果当前条件是MMN，则发送trigger "32"
            fwrite(Obj,trig32);
        end
        if cond_idx==3% 如果当前条件是AHER，则发送trigger "33"
            fwrite(Obj,trig33);
        end

        pause(1);
        Beeper(1000, 1, 1); % audio播放结束的提示音

        pause(questionnaire_time) % 20秒钟填写digitalized SAM

        set(txt1,'visible','off');
        set(txt2,'visible','on');
    end
end
Beeper(1000, 1, 1);

fclose(scom);
fclose(Obj);
close all;
toc

%% 关闭串口
fclose(Obj);
clear global Obj
```
