## Supplementary material for "Auditory Stimulus Information Entropy Modulates Inter-Brain Synchronization: Evidence from Wireless EEG Hyperscanning": program code: send_trigger.m

```
function send_trigger(ifsend, trigger)
% ifsend: 0 = not send; 1 = send.

if ifsend == 1
    global Obj;
    hex_trigger = dec2hex(trigger);
    if trigger < 16
        hex_trigger = [0,hex_trigger];
    end
    trig = [hex_trigger; '55'; '66';'0D'];
    trig_left = hex2dec(trig);

    fwrite(Obj, trig_left);
end

end
```
