## Supplementary material for "Auditory Stimulus Information Entropy Modulates Inter-Brain Synchronization: Evidence from Wireless EEG Hyperscanning": program code: song2_Trigger.m

```
% for times = 1:5
%     song2_test;
% end

% clc;clear;C:\Users\CBHguest\Desktop\Liao\SCR_Brain_Music\Experiment_2
%% 記得改回180
% play_time = 20 + 1; % unit: seconds 
%% MIDI
% devinfo = mididevinfo;
device = mididevice('Output',3); % 電子鋼琴
% device = mididevice('Output',1);
msgs = midimsg('ProgramChange',1,11);
midisend(device,msgs);
peroid1 = 1/6;
tic;
flash_time1 = 1;
k=1;
p=0;
while 1
    t=toc;
    if t>flash_time1
        if flash_time1 == 1 % 开始时发送1
            fwrite(Obj,trig6);  % 发送表示DW条件开始的trigger
        end
        clear msg;
        switch k
            case 1
                msg(1) = midimsg('NoteOn',1,33+p,40,0);
                msg(2) = midimsg('NoteOff',1,33+p,40,0.1);
                fwrite(Obj,trig7); % 每个音符都发送2
                midisend(device,msg);
            case 2
                msg(1) = midimsg('NoteOn',1,p+40,40,0);
                msg(2) = midimsg('NoteOff',1,p+40,40,0.1);
                fwrite(Obj,trig7);
                midisend(device,msg);
            case 3
                msg(1) = midimsg('NoteOn',1,48+p,40,0);
                msg(2) = midimsg('NoteOff',1,48+p,40,0.1);
                fwrite(Obj,trig7);
                midisend(device,msg);
            case 4
                msg(1) = midimsg('NoteOn',1,40+p,40,0);
                msg(2) = midimsg('NoteOff',1,40+p,40,0.1);
                fwrite(Obj,trig7);
                midisend(device,msg);
            case 5
                msg(1) = midimsg('NoteOn',1,48+p,40,0);
                msg(2) = midimsg('NoteOff',1,48+p,40,0.1);
                fwrite(Obj,trig7);
                midisend(device,msg);
            case 6
                msg(1) = midimsg('NoteOn',1,40+p,40,0);
                msg(2) = midimsg('NoteOff',1,40+p,40,0.1);
                fwrite(Obj,trig7);
                midisend(device,msg);
            case 7
                msg(1) = midimsg('NoteOn',1,48+p,40,0);
                msg(2) = midimsg('NoteOff',1,48+p,40,0.1);
                fwrite(Obj,trig7);
                midisend(device,msg);
            case 8
                msg(1) = midimsg('NoteOn',1,40+p,40,0);
                msg(2) = midimsg('NoteOff',1,40+p,40,0.08);
                fwrite(Obj,trig7);
                midisend(device,msg);
            case 9
                msg(1) = midimsg('NoteOn',1,69+p,60,0);
                msg(2) = midimsg('NoteOff',1,69+p,60,0.11);
                fwrite(Obj,trig7);
                midisend(device,msg);
            case 10
                msg(1) = midimsg('NoteOn',1,p+69,60,0);
                msg(2) = midimsg('NoteOff',1,p+69,60,0.08);
                msg(3) = midimsg('NoteOn',1,33+p,40,0);
                msg(4) = midimsg('NoteOff',1,33+p,40,0.08);
                fwrite(Obj,trig7);
                midisend(device,msg);
            case 11
                msg(1) = midimsg('NoteOn',1,71+p,60,0);
                msg(2) = midimsg('NoteOff',1,71+p,60,0.1);
                fwrite(Obj,trig7);
                midisend(device,msg);
            case 12
                msg(1) = midimsg('NoteOn',1,71+p,60,0);
                msg(2) = midimsg('NoteOff',1,71+p,60,0.08);
                fwrite(Obj,trig7);
                midisend(device,msg);
            case 13
                msg(1) = midimsg('NoteOn',1,72+p,60,0);
                msg(2) = midimsg('NoteOff',1,72+p,60,0.1);
                fwrite(Obj,trig7);
                midisend(device,msg);
            case 14
                msg(1) = midimsg('NoteOn',1,p+72,60,0);
                msg(2) = midimsg('NoteOff',1,p+72,60,0.08);
                fwrite(Obj,trig7);
                midisend(device,msg);
            case 15
                msg(1) = midimsg('NoteOn',1,71+p,60,0);
                msg(2) = midimsg('NoteOff',1,71+p,60,0.1);
                fwrite(Obj,trig7);
                midisend(device,msg);
            case 16
                msg(1) = midimsg('NoteOn',1,71+p,60,0);
                msg(2) = midimsg('NoteOff',1,71+p,60,0.08);
                fwrite(Obj,trig7);
                midisend(device,msg);
            case 17
                msg(1) = midimsg('NoteOn',1,69+p,60,0);
                msg(2) = midimsg('NoteOff',1,69+p,60,0.1);
                fwrite(Obj,trig7);
                midisend(device,msg);
            case 18
                msg(1) = midimsg('NoteOn',1,p+69,60,0);
                msg(2) = midimsg('NoteOff',1,p+69,60,0.08);
                msg(3) = midimsg('NoteOn',1,33+p,40,0);
                msg(4) = midimsg('NoteOff',1,33+p,40,0.08);
                fwrite(Obj,trig7);
                midisend(device,msg);
            case 19
                msg(1) = midimsg('NoteOn',1,64+p,60,0);
                msg(2) = midimsg('NoteOff',1,64+p,60,0.1);
                fwrite(Obj,trig7);
                midisend(device,msg);
            case 20
                msg(1) = midimsg('NoteOn',1,64+p,60,0);
                msg(2) = midimsg('NoteOff',1,64+p,60,0.08);
                fwrite(Obj,trig7);
                midisend(device,msg);
            case 21
                msg(1) = midimsg('NoteOn',1,60+p,60,0);
                msg(2) = midimsg('NoteOff',1,60+p,60,0.1);
                fwrite(Obj,trig7);
                midisend(device,msg);
            case 22
                msg(1) = midimsg('NoteOn',1,p+60,60,0);
                msg(2) = midimsg('NoteOff',1,p+60,60,0.08);
                fwrite(Obj,trig7);
                midisend(device,msg);
            case 23
                msg(1) = midimsg('NoteOn',1,57+p,60,0);
                msg(2) = midimsg('NoteOff',1,57+p,60,0.1);
                fwrite(Obj,trig7);
                midisend(device,msg);
            case 24
                msg(1) = midimsg('NoteOn',1,57+p,60,0);
                msg(2) = midimsg('NoteOff',1,57+p,60,0.08);
                fwrite(Obj,trig7);
                midisend(device,msg);
            case 25
                msg(1) = midimsg('NoteOn',1,67+p,60,0);
                msg(2) = midimsg('NoteOff',1,67+p,60,0.1);
                fwrite(Obj,trig7);
                midisend(device,msg);
            case 26
                msg(1) = midimsg('NoteOn',1,p+67,60,0);
                msg(2) = midimsg('NoteOff',1,p+67,60,0.08);
                msg(3) = midimsg('NoteOn',1,38+p,40,0);
                msg(4) = midimsg('NoteOff',1,38+p,40,0.08);
                fwrite(Obj,trig7);
                midisend(device,msg);
            case 27
                msg(1) = midimsg('NoteOn',1,65+p,60,0);
                msg(2) = midimsg('NoteOff',1,65+p,60,0.1);
                fwrite(Obj,trig7);
                midisend(device,msg);
            case 28
                msg(1) = midimsg('NoteOn',1,65+p,60,0);
                msg(2) = midimsg('NoteOff',1,65+p,60,0.08);
                fwrite(Obj,trig7);
                midisend(device,msg);
            case 29
                msg(1) = midimsg('NoteOn',1,64+p,60,0);
                msg(2) = midimsg('NoteOff',1,64+p,60,0.1);
                fwrite(Obj,trig7);
                midisend(device,msg);
            case 30
                msg(1) = midimsg('NoteOn',1,p+65,60,0);
                msg(2) = midimsg('NoteOff',1,p+65,60,0.08);
                fwrite(Obj,trig7);
                midisend(device,msg);
            case 31
                msg(1) = midimsg('NoteOn',1,p+67,60,0);
                msg(2) = midimsg('NoteOff',1,p+67,60,0.1);
                fwrite(Obj,trig7);
                midisend(device,msg);
            case 32
                msg(1) = midimsg('NoteOn',1,p+65,60,0);
                msg(2) = midimsg('NoteOff',1,p+65,60,0.12);
                msg(3) = midimsg('NoteOn',1,38+p,40,0);
                msg(4) = midimsg('NoteOff',1,38+p,40,0.12);
                fwrite(Obj,trig7);
                midisend(device,msg);
            case 33
                msg(1) = midimsg('NoteOn',1,p+31,40,0);
                msg(2) = midimsg('NoteOff',1,p+31,40,0.1);
                fwrite(Obj,trig7);
                midisend(device,msg);
            case 34
                msg(1) = midimsg('NoteOn',1,p+65,60,0);
                msg(2) = midimsg('NoteOff',1,p+65,60,0.1);
                fwrite(Obj,trig7);
                midisend(device,msg);
            case 35
                msg(1) = midimsg('NoteOn',1,p+65,60,0);
                msg(2) = midimsg('NoteOff',1,p+65,60,0.08);
                fwrite(Obj,trig7);
                midisend(device,msg);
            case 36
                msg(1) = midimsg('NoteOn',1,p+67,60,0);
                msg(2) = midimsg('NoteOff',1,p+67,60,0.1);
                fwrite(Obj,trig7);
                midisend(device,msg);
            case 37
                msg(1) = midimsg('NoteOn',1,p+67,60,0);
                msg(2) = midimsg('NoteOff',1,p+67,60,0.08);
                fwrite(Obj,trig7);
                midisend(device,msg);
            case 38
                msg(1) = midimsg('NoteOn',1,p+69,40,0);
                msg(2) = midimsg('NoteOff',1,p+69,40,0.1);
                fwrite(Obj,trig7);
                midisend(device,msg);
            case 39
                msg(1) = midimsg('NoteOn',1,p+69,60,0);
                msg(2) = midimsg('NoteOff',1,p+69,60,0.08);
                fwrite(Obj,trig7);
                midisend(device,msg);
            case 40
                msg(1) = midimsg('NoteOn',1,p+71,40,0);
                msg(2) = midimsg('NoteOff',1,p+71,40,0.1);
                fwrite(Obj,trig7);
                midisend(device,msg);
            case 41
                msg(1) = midimsg('NoteOn',1,p+71,60,0);
                msg(2) = midimsg('NoteOff',1,p+71,60,0.08);
                msg(3) = midimsg('NoteOn',1,p+31,40,0);
                msg(4) = midimsg('NoteOff',1,p+31,40,0.08);
                fwrite(Obj,trig7);
                midisend(device,msg);
            case 42
                msg(1) = midimsg('NoteOn',1,p+67,40,0);
                msg(2) = midimsg('NoteOff',1,p+67,40,0.1);
                fwrite(Obj,trig7);
                midisend(device,msg);
            case 43
                msg(1) = midimsg('NoteOn',1,p+67,60,0);
                msg(2) = midimsg('NoteOff',1,p+67,60,0.08);
                fwrite(Obj,trig7);
                midisend(device,msg);
            case 44
                msg(1) = midimsg('NoteOn',1,p+62,60,0);
                msg(2) = midimsg('NoteOff',1,p+62,60,0.1);
                fwrite(Obj,trig7);
                midisend(device,msg);
            case 45
                msg(1) = midimsg('NoteOn',1,p+62,60,0);
                msg(2) = midimsg('NoteOff',1,p+62,60,0.08);
                fwrite(Obj,trig7);
                midisend(device,msg);
            case 46
                msg(1) = midimsg('NoteOn',1,p+65,40,0);
                msg(2) = midimsg('NoteOff',1,p+65,40,0.1);
                fwrite(Obj,trig7);
                midisend(device,msg);
            case 47
                msg(1) = midimsg('NoteOn',1,p+65,60,0);
                msg(2) = midimsg('NoteOff',1,p+65,60,0.08);
                msg(3) = midimsg('NoteOn',1,p+36,40,0);
                msg(4) = midimsg('NoteOff',1,p+36,40,0.08);
                fwrite(Obj,trig7);
                midisend(device,msg);
            case 48
                msg(1) = midimsg('NoteOn',1,p+64,60,0);
                msg(2) = midimsg('NoteOff',1,p+64,60,0.1);
                fwrite(Obj,trig7);
                midisend(device,msg);
            case 49
                msg(1) = midimsg('NoteOn',1,p+64,60,0);
                msg(2) = midimsg('NoteOff',1,p+64,60,0.08);
                fwrite(Obj,trig7);
                midisend(device,msg);
            case 50
                msg(1) = midimsg('NoteOn',1,p+62,40,0);
                msg(2) = midimsg('NoteOff',1,p+62,40,0.1);
                fwrite(Obj,trig7);
                midisend(device,msg);
            case 51
                msg(1) = midimsg('NoteOn',1,p+64,60,0);
                msg(2) = midimsg('NoteOff',1,p+64,60,0.08);
                fwrite(Obj,trig7);
                midisend(device,msg);
            case 52
                msg(1) = midimsg('NoteOn',1,p+65,60,0);
                msg(2) = midimsg('NoteOff',1,p+65,60,0.1);
                fwrite(Obj,trig7);
                midisend(device,msg);
            case 53
                msg(1) = midimsg('NoteOn',1,p+64,60,0);
                msg(2) = midimsg('NoteOff',1,p+64,60,0.11);
                msg(3) = midimsg('NoteOn',1,p+40,40,0);
                msg(4) = midimsg('NoteOff',1,p+40,40,0.11);
                fwrite(Obj,trig7);
                midisend(device,msg);
            case 54
                msg(1) = midimsg('NoteOn',1,p+76,60,0);
                msg(2) = midimsg('NoteOff',1,p+76,60,0.13);
                fwrite(Obj,trig7);
                midisend(device,msg);
        end
        k=k+1;
        if k>54
            p=p+1;
            k=1;
        end
        flash_time1 = flash_time1 + peroid1;
    end
    if flash_time1 > play_time
        trig36 = hex2dec(['24';'55';'66';'0D']); 
        fwrite(Obj,trig36); % 发送表示DW条件结束的trigger
        break
        % return;
    end
end
% send_trigger(1, 36);
% trig36 = hex2dec(['24';'55';'66';'0D']); 
% fwrite(Obj,trig36); % 发送表示DW条件结束的trigger
%% 关闭串口
% fclose(Obj);
% clear global Obj
```
